# Pancreatic islet chromatin accessibility and conformation defines distal enhancer networks of type 2 diabetes risk

**DOI:** 10.1101/299388

**Authors:** William W Greenwald, Joshua Chiou, Jian Yan, Yunjiang Qiu, Ning Dai, Allen Wang, Naoki Nariai, Anthony Aylward, Jee Yun Han, Nikita Kadakia, Laura Barrufet, Mei-Lin Okino, Frauke Drees, Nicholas Vinckier, Liliana Minichiello, David Gorkin, Joseph Avruch, Kelly Frazer, Maike Sander, Bing Ren, Kyle J Gaulton

## Abstract

The gene targets of enhancer activity in pancreatic islets are largely unknown, impeding discovery of islet regulatory networks involved in type 2 diabetes (T2D) risk. We mapped chromatin state, accessibility and conformation using ChIP-seq, ATAC-seq and Hi-C in human pancreatic islets, which we integrated with T2D genetic fine-mapping and islet expression QTL data. Active islet regulatory elements preferentially interacted with other active elements, often at distances over 1MB, and we identified target genes for thousands of distal islet enhancers. A third of T2D risk signals mapped in islet enhancers, and target genes regulated by these signals were specifically involved in processes related to protein transport and secretion. Among implicated target genes of T2D islet enhancer signals with no prior known role in islet function, we demonstrated that reduced *IGF2BP2* activity in mouse islets leads to impaired glucose-stimulated insulin secretion. These results link distal islet enhancer regulation of protein secretion and transport to genetic risk of T2D, and highlight the utility of high-throughput chromatin conformation maps to uncover the gene regulatory networks of complex disease.

## Introduction

Genetic risk of type 2 diabetes (T2D) is largely mediated through variants affecting transcriptional regulatory activity in pancreatic islets^1–7^. Genetic fine-mapping combined with epigenomic annotation data can identify causal variants at T2D risk loci mapping in islet regulatory elements^1,2^. The gene targets of islet regulatory elements, however, are largely unknown, impeding discovery of disease-relevant gene networks perturbed by risk variants and novel therapeutic avenues. The spatial organization of chromatin plays a critical role in tissue-specific gene regulation, and recently developed high-throughput techniques such as Hi-C identify physical relationships between genomic regions in human tissues genome-wide^8–10^. Tissue-specific maps of chromatin organization can identify candidate target genes of distal regulatory elements and inform the molecular mechanisms of disease risk variants^9^.

Here, we defined the spatial organization of transcriptional regulatory elements in primary pancreatic islets, through which we mapped genetic effects on islet gene expression and T2D risk. Islet active regulatory elements preferentially interacted with other active elements, in many cases over 1MB, and we identified putative distal target genes for thousands of islet enhancers. A third of known T2D risk signals had likely causal variants in islet enhancers, and target genes of these signals were strongly enriched for processes related to protein secretion and transport. Among target genes with no previously known role in islet function, we demonstrated that reduced activity of *IGF2BP2* in mouse islets leads to impaired glucose-stimulated insulin secretion. Together our results define distal regulatory programs in islets through which we link islet enhancer regulation of protein transport and secretion to T2D risk.

## Results

We first defined islet accessible chromatin using ATAC-seq^11^ generated from four pancreatic islet samples (**Table S1**). Accessible chromatin signal was highly concordant across all samples (Pearson r^2^>.8) (**Figure S1**). We called sites for each sample separately using MACS2^12^, and merged sites to create a combined set of 105,734 islet accessible chromatin sites. We then collected published ChIP-seq data of histone modification and transcription factor binding in primary islets^5,13^ and called chromatin states using ChromHMM^14^ (**Figure S1**). Accessible chromatin predominantly mapped within active enhancer and promoter states (**Figure 1A**). We functionally annotated islet accessible chromatin by using chromatin states to define active enhancers and promoters, as well as other classes of islet regulatory elements (**Table S2**). We identified 44,860 active enhancers which, in line with previous reports^4,15^, were distal to promoters, more tissue-specific, and preferentially harbored motifs for FOXA, RFX, NEUROD and other islet transcription factors (**Figure S1, Table S3**). These results define active enhancers and other classes of regulatory elements in pancreatic islets.

**Figure 1.**
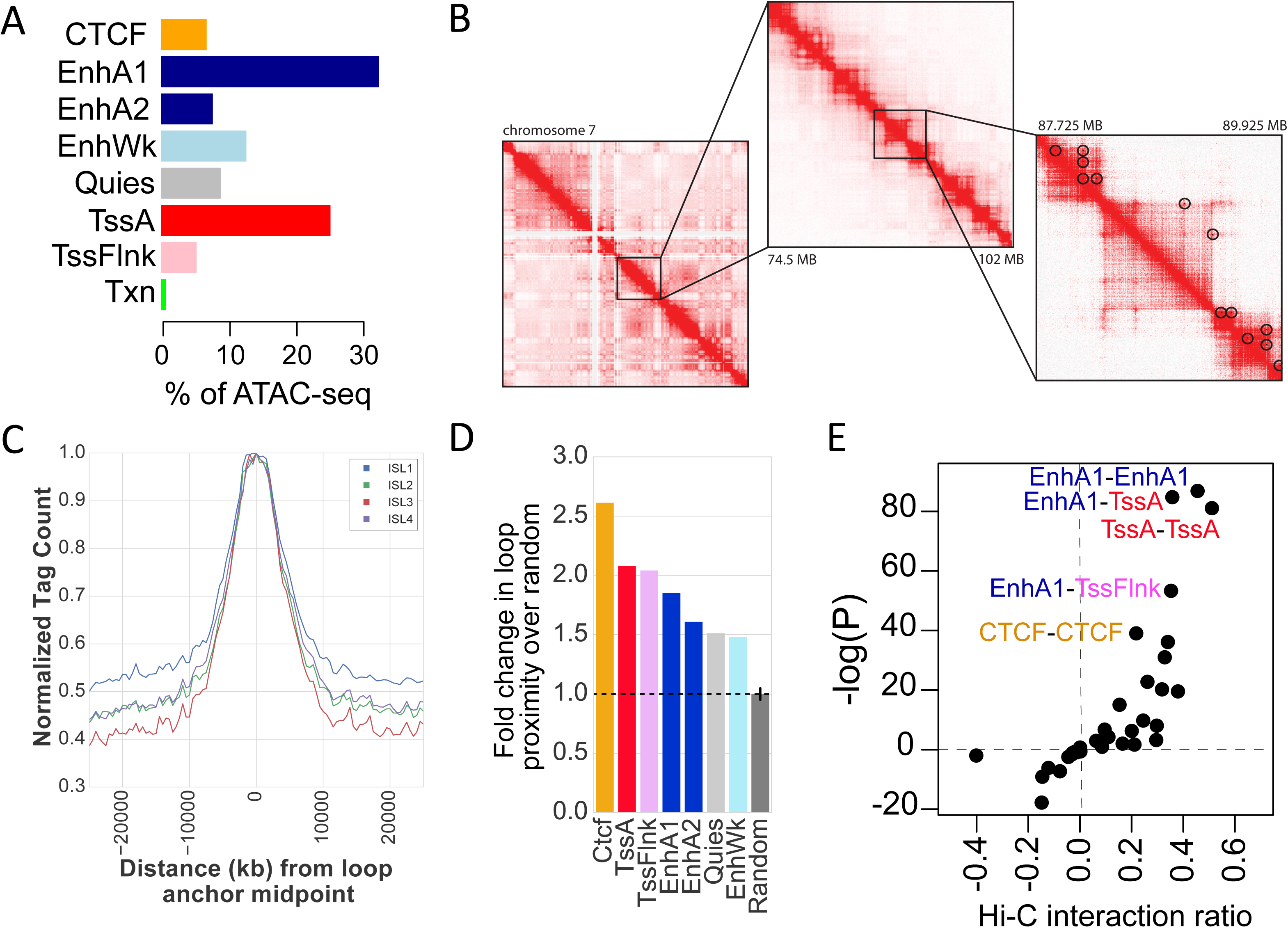
Chromatin accessibility and conformation in pancreatic islets. (A) Islet accessible chromatin signal mapped predominantly within active islet chromatin states. (B) Chromatin looping from *in situ* Hi-C assays of three pancreatic islet samples at entire chromosome (left), 25MB (middle) and 2MB (right) resolution on chromosome 7. Black circles on the right panel represent statistically significant loop calls. (C) Accessible chromatin signal from four islet samples (ISL1-4) was distributed around chromatin loop anchor midpoints (D) Islet chromatin loop anchors were enriched for islet CTCF binding sites, active enhancers, and active promoters compared to random sites. Values represent fold change and SD. (E) Islet chromatin loops were most enriched for interactions between islet active enhancers and active promoter elements, and between CTCF binding sites.

Defining the target genes of enhancers has been challenging as they frequently control non-adjacent genes over large genomic distances through chromatin looping^16^. The spatial organization of chromatin in pancreatic islets is unknown, and we therefore identified physical interactions between genomic regions in islets. We performed genome-wide chromatin conformation capture using *in situ* Hi-C^8,17^ in three islet samples, two of which were sequenced to a depth of >1 billion reads (**Table S1**). Contact matrices from islet Hi-C assays were highly concordant across samples (**Figure S2**). We called chromatin loops at 5kb, 10kb, and 25kb resolution with HICCUPS^8^ using reads from each sample individually, as well as with reads pooled from all three samples (**Figure 1B**). We then merged loops across samples where both anchors overlapped at 20kb resolution (**see Methods**), resulting in a combined set of 11,924 islet Hi-C loops (**Table S4**). The median distance between loop anchor midpoints was 255kb, and nearly 10% were over 1MB in size (**Figure S2**). This established a map of chromatin loops in human pancreatic islets.

We determined the relationship between islet regulatory element activity and chromatin looping. Islet accessible chromatin signal was largely localized to islet loop anchors, with the strongest signal at anchor midpoints (**Figure 1C**). Nearly half of all islet regulatory elements were proximal to an anchor (48.7%), and regulatory sites most enriched (empirical P<1.5×10^−4^) for chromatin loop anchors included CTCF binding sites (2.61-fold), active promoters (2.08-fold), and active enhancers (1.85-fold) (**Figure 1D**). We further mapped the relationship between islet regulatory sites connected by loop anchors. The most significantly enriched anchor interactions were between active enhancer and promoter sites (EnhA1-TssA OR=1.28, P=1.53×10^−37^; EnhA1-EnhA1 OR=1.37, P=1.87×10^−38^; TssA-TssA OR= 1.42, P=6.15×10^−36^). We also observed strong enrichment for interactions between CTCF binding sites (CTCF-CTCF OR=1.16; P=1.1×10^−17^) (**Figure 1E**). These results demonstrate that islet chromatin loops are prominently enriched for active regulatory sites in addition to CTCF binding sites.

We next used chromatin loops to annotate distal relationships between islet enhancers and potential target genes genome-wide (**see Methods**). Over 40% (18,240) of islet active enhancer elements interacted with at least one gene promoter region, and on average, these enhancers interacted with 2 gene promoter regions (**Figure S2, Table S5**). Conversely, the promoter regions of 8,448 genes had at least one loop to an enhancer element (**Figure 2A, Table S6**). Of these 8,448 genes, 1,157 had more than two independent chromatin loops to enhancer elements. Genes with multiple loops were strongly enriched for processes related to transcription factor activity and gene regulation, protein transport, and insulin signaling (**Table S7**). Among highly-looped genes were also numerous critical for islet function, such as *INS*, *ISL1*, *FOXA2*, *NKX6.1*, and *MAFB* (**Table S5**). For example, there were four distinct loops between active enhancers and *MAFB*, including several loops to enhancers over 1 MB distal (**Figure 2B**). These results define candidate distal target genes of enhancer elements in islets.

**Figure 2.**
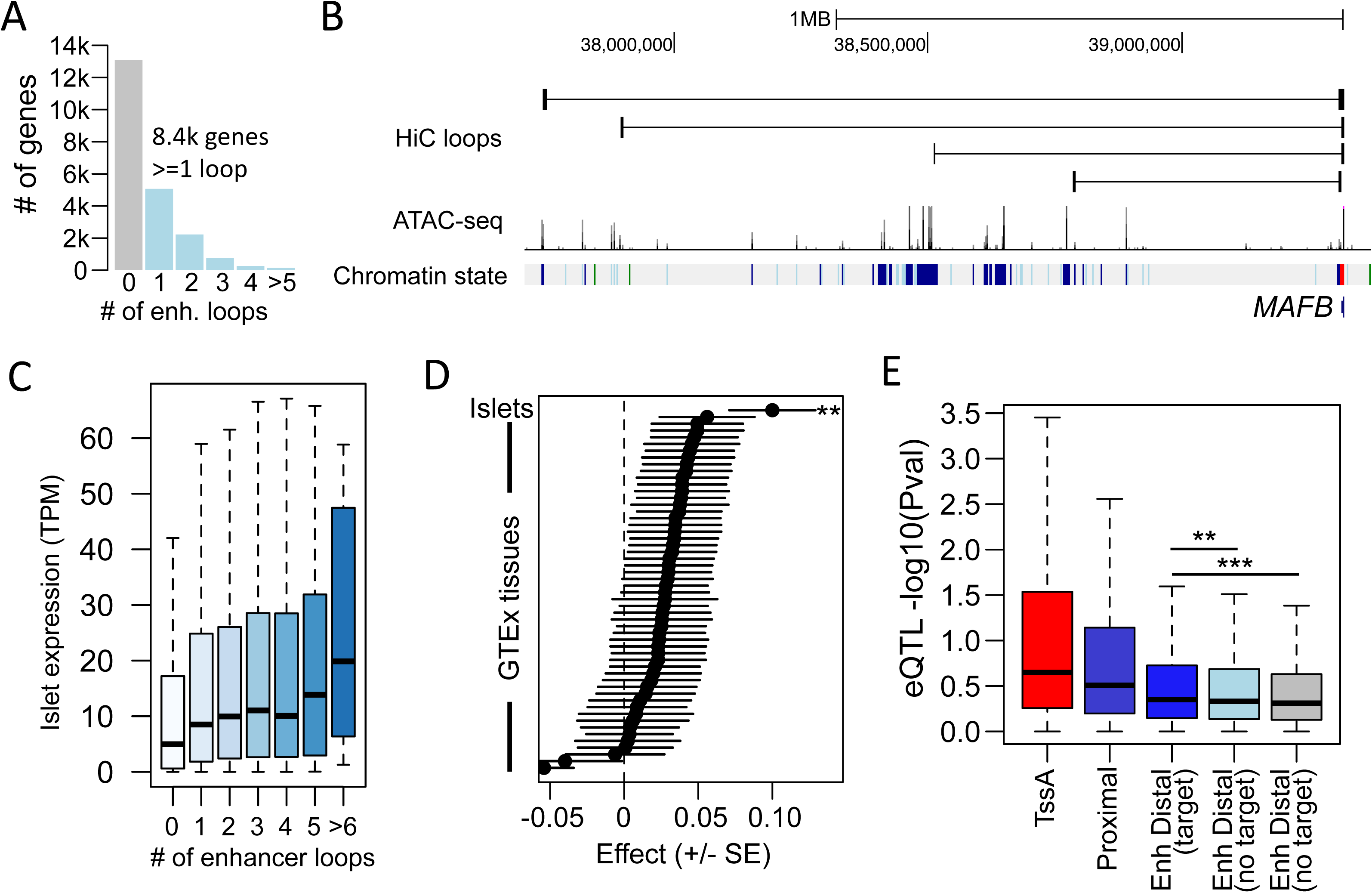
Islet enhancer regulation of distal target gene expression. (A) The promoter regions of 8.4k genes had at least one chromatin loop to an islet enhancer element. (B) Multiple islet enhancers formed chromatin loops with the *MAFB* promoter region including several over 1MB. (C) Genes with increasing numbers of chromatin loops to islet enhancers had increased expression level in islets, with the highest expression among genes with 6 or more interactions (D) The number of chromatin loops to islet enhancers was a significant predictor of islet gene expression but not 53 other tissues in GTEx. Values represent effect size and SE. ^⋆⋆^P<.001 (E) Genetic variants in distal islet enhancer elements had stronger evidence for islet expression QTLs with genes in chromatin loops (blue) than genes with no loop (grey), even when matched based on distance (light blue). ^⋆⋆^P<.001, ^⋆⋆⋆^P<.0001

We examined the relationship between active enhancer interactions and target gene expression level by using RNA-seq data from pancreatic islet samples^18^ and 53 tissues in GTEx release v7 data^19^. A significantly higher proportion of genes expressed in islets had at least one enhancer loop compared to non-islet expressed genes (ln(TPM)>1; expr=.48, non-expr=.30, Chi-square P=2.2×10^−16^). Genes with increasing numbers of enhancer loops had, on average, higher expression level in islets (ρ=.15, P=2.2×10^−16^), with the highest expression among genes with 6 or more loops (median=19.8 TPM) (**Figure 2C**). The number of islet enhancer interactions was also a significant predictor of expression level in islets (β=.10, P=6.4×10^−4^), and not of relative expression level in any of the other 53 tissues in GTEx (all P>.05) (**Figure 2D**). We measured the relative expression level of genes in islets and 53 GTEx tissues normalized across all tissues (**see Methods**), and again observed a significant relationship between enhancer loops and relative expression level in islets and no other tissues (**Figure S2**). These results suggest distal islet enhancer chromatin loops are correlated with islet-specific gene expression patterns.

We next determined the effects of genetic variants in islet enhancers on target gene regulation. We generated expression quantitative trait loci (eQTL) data in 230 islet RNA-seq samples by combining summary statistics from two published studies through meta-analysis^7,18^ (**see Methods**). We identified variants overlapping classes of islet regulatory elements genome-wide. We then quantified the eQTL association of these variants to target genes determined from their proximity to nearby genes and from chromatin loops (**see Methods**). As expected, we observed the strongest eQTL evidence for active promoter and enhancer variants proximal to genes (TssA median –log10(P)=.65, EnhA proximal median –log10(P)=.50) (**Figure 2E**). For variants in distal enhancers, we observed stronger evidence for islet eQTL association among genes in loops relative to non-loop genes (EnhA interacting median=.35, EnhA non-interacting median=.31, Wilcox P=2.2×10^−16^), even when matching based on gene distance to the enhancer (Wilcox P=2.9×10^−4^) (**Figure 2E**). These results suggest that genetic variants in distal islet enhancer elements are preferentially correlated with the expression level of genes within chromatin loops.

Genetic variants at T2D risk loci are enriched for islet regulatory elements^1,2,4,5^, but the effects of variants in regulatory elements on T2D risk in the context of chromatin looping is unknown. We determined the effects of variants in islet regulatory elements and chromatin loops on T2D risk using association data of 6.1M common (MAF>.05) variants with fgwas and LD-score regression^20,21^. We observed strongest enrichment of variants in active regulatory elements, most notably in active enhancers (EnhA1 fgwas ln(enrich)=3.9, LD-score Z=3.1) (**Figure 3A, Figure S3**). The effects of variants in active enhancer and promoter elements on T2D risk were more pronounced among those in chromatin loops (EnhA1 fgwas ln(enrich)=4.38, LD-score Z=3.1; TssA fgwas ln(enrich)=3.03, LD-score Z=0.86) (**Figure 3B, Figure S3**). Conversely, variants in other islet elements such as flanking promoters and weak enhancers and were more enriched outside of loops (**Figure 3B, Figure S3**). To determine the inter-dependence of these effects, we jointly modelled variants in islet regulatory elements on T2D risk, while also including variants in GENCODE coding exons and UTRs. In a joint model, we observed enrichment of variants in islet active enhancer elements (EnhA1 ln(enrich)=4.04), in addition to flanking promoters (TssFlnk ln(enrich)=3.77) and coding exons (CDS ln(enrich)=2.34) (**Figure S3**). These results demonstrate genome-wide enrichment of variants in islet active regulatory elements within chromatin loops for T2D risk.

**Figure 3.**
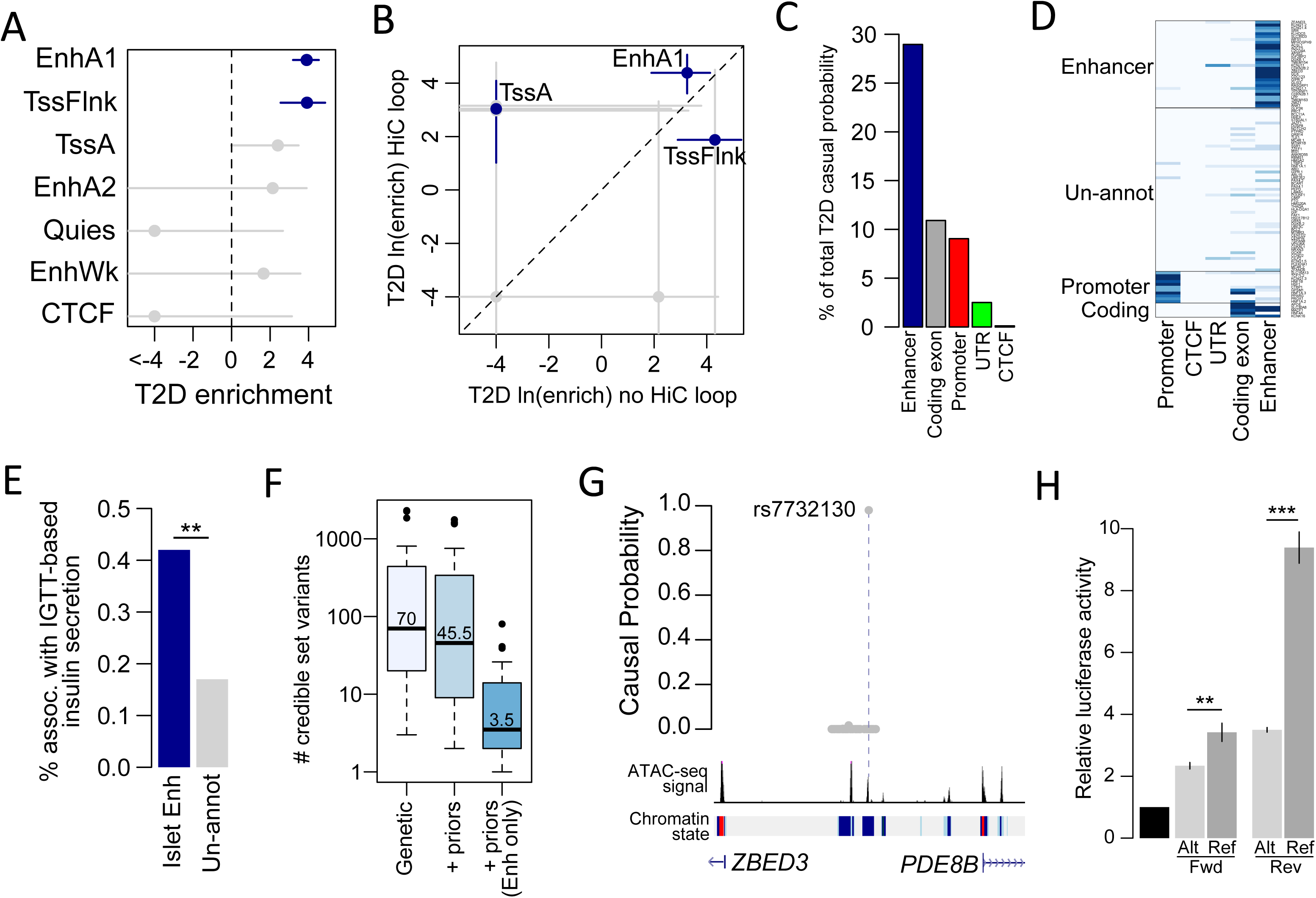
Type 2 diabetes risk signals map in islet enhancers. (A) Genetic variants in islet active regulatory elements genome-wide were enriched for T2D risk, with strongest enrichment in active enhancer elements. Values represent log enrichment and 95% CI. (B) The effects of variants in active enhancer and promoter elements on T2D risk were stronger among those in chromatin loops, whereas other elements were enriched for T2D outside of loops. Values represent log enrichment and 95% CI. (C) Over 30% of the total causal probability across 107 known T2D risk signals mapped in islet enhancer elements. (D) Clustering of known T2D signals based on islet and coding annotations identified 30 signals with likely causal variants in islet enhancers. (E) A significantly higher percentage of T2D islet enhancer signals were associated with IGTT-based insulin secretion phenotypes than un-annotated T2D signals. ^⋆⋆^P<.001 (F) Number of variants in the 99% credible sets for the 30 T2D islet enhancer signals based on genetic fine-mapping alone (genetic), genetic fine-mapping including functional priors (+priors) (G) T2D causal variant rs7732130 at the *ZBED3* locus mapped in an islet active enhancer and chromatin loop, and had (H) allelic effects on enhancer activity in the islet cell line MIN6 (N=3). Values represent fold change and SD. ^⋆⋆^P<.001, ^⋆⋆⋆^P<.0001

To identify T2D risk signals mapping in islet enhancers, we used effects from the joint enrichment model as priors on the causal evidence (PPA) for both variants at known T2D loci and in windows genome-wide^1,2,21^ (**see Methods**). Among 107 known risk signals, variants in islet enhancers accounted for almost a third (29%) of the total probability mass (**Table S8, Figure 3C**). We clustered known risk signals based on annotations at candidate causal variants (**see Methods**) and identified 30 signals where the causal variant was likely in an islet enhancer (**Figure 3D**). The 30 T2D islet enhancer signals were associated with IGTT-based insulin secretion phenotypes significantly more than un-annotated signals (Enh=42%, un-annot=17%, Chi-square P=1×10^−7^), supporting a role in islet function^22^ (**Figure 3E**). Fine-mapping including functional priors improved resolution of causal variants at the 30 T2D islet enhancer signals (avg. 3.5 enh variants) (Figure 3F), and at 6 signals resolved a single causal islet enhancer variant (**Table S8**). One example is previously unreported variant rs7732130 (*ZBED3;* PPA=98%) in a chromatin loop and which has allelic effects on islet enhancer activity (T-test Fwd P=3.7×10^−3^, Rev P=6.8×10^−6^) (**Figure 3G, 3H**). Outside of known loci, we identified an additional 131 1MB windows genome-wide harboring putative T2D enhancer variants (**Figure S3, Table S9; see Methods**). These results identify a large number of known and putative T2D risk signals with causal variants in islet enhancers.

A large percentage of T2D risk signals map in islet enhancers, and the gene targets of these signals are largely unknown. We defined candidate target genes based on gene promoter regions in chromatin loops with, or in proximity to, T2D enhancer signals (**Figure 3D, see Methods**). T2D enhancer signals had on average 2 target genes (**Figure 4A, 4B, Table S10**), a large reduction in candidate gene numbers obtained when using a 1MB window (avg.=18 genes) or TAD definitions (avg.=7 genes) (**Figure 4A**). Target genes were enriched in gene sets related to protein transport and secretion, potassium ion transport, vesicles and vesicle membranes, and endoplasmic reticulum (FDR<.2) (**Figure 4B, Table S11**). Target genes also included multiple involved in MODY and other monogenic and syndromic diabetes (*ABCC8*, *KCNJ11*, *GCK*, *INS*, *GLIS3*, *WFS1*) (**Figure S4**). Conversely, non-target genes within 1MB of these same 30 signals were enriched for gene sets related to stress-response and other processes (FDR<.2), suggesting regulatory programs potentially activated in other cellular states (**Table S11, see Methods**). At several loci, loops implicated target genes highly distal (>500kb) to T2D enhancer variants; for example, multiple *KCNQ1* signals interacted with *INS/IGF2* over 700kb distal, and *ZMIZ1* interacted with *POLR3A* over 1MB distal (**Figure S4**). These results define putative target genes of T2D enhancer signals involved in protein transport and secretion and monogenic diabetes.

**Figure 4.**
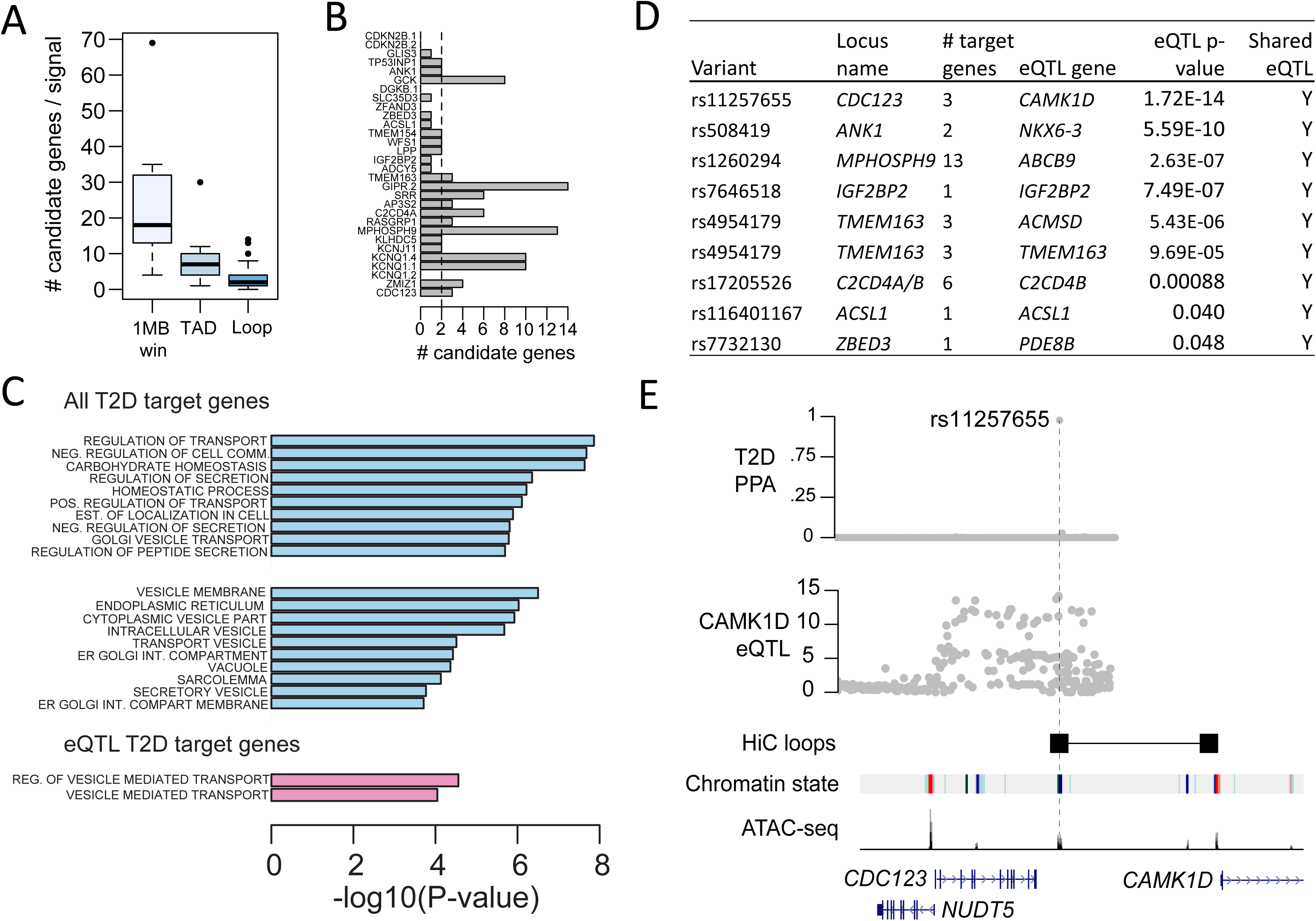
Target genes of type 2 diabetes islet enhancer signals are involved in protein secretion and transport. (A) Prioritizing target genes using chromatin loops and proximity greatly reduces the number of candidate genes over using a 1MB window (avg=18) or topologically associated domain (TAD) boundaries (avg=7). (B) T2D islet enhancer signals formed chromatin loops with, or were in proximity to, an average of 2 target genes. (C) Target genes of T2D enhancer signals were strongly enriched for biological processes related to protein secretion, protein transport, vesicles and vesicle membranes, and endoplasmic reticulum (FDR<.2) (top), and target genes with islet eQTL evidence were specifically enriched for vesicle-mediated transport (FDR<.2) (bottom). (D) Target genes with islet eQTLs to T2D islet enhancer signals (corrected eQTL P<.05; p-values reported in table are uncorrected) and evidence that T2D and eQTL signals are co-localized. (E) At the *CDC123/CAMK1D* locus T2D islet enhancer variant rs11257655 was in a chromatin loop to the *CAMK1D* promoter and an islet eQTL for *CAMK1D* expression. Probabilities (PPA) that variants are causal for T2D risk (top) and variant association (-log10 P) with islet expression level of *CAMK1D* (middle).

We then further identified target genes regulated by T2D enhancer signals using islet eQTL data. At each signal, we tested the most likely casual enhancer variant for eQTL association to each target gene correcting for the total number of target genes (**see Methods**). For genes with eQTL evidence (P<.05), we further confirmed eQTL and T2D signals were unlikely to be driven by distinct causal variants using Bayesian co-localization (**see Methods**). Target genes showed evidence of islet eQTLs with 8 known T2D islet enhancer signals (P<.05) including *CAMK1D*, *ABCB9*, *C2CD4B*, and *IGF2BP2* (**Figure 4D, Table S12**). For example, known T2D variant rs11257655 is in an islet active enhancer element that loops to the *CAMK1D* promoter and is an islet eQTL for *CAMK1D* expression^23^ (**Figure 4E**). At the 131 putative T2D enhancer signals, we identified 12 additional target genes with evidence for eQTLs to T2D variants (P<.05) such as *FADS1*, *VEGFA*, *SNX32* and *SCRN2* (**Table S12**). Among the 21 directly regulated genes, nearly a third have not been identified as significant islet eQTLs in previous studies^7,18,41^. Target genes with islet eQTLs to known and putative T2D enhancer signals were specifically enriched for genes involved in vesicle-mediated transport (FDR<.2) (**Figure 4C, Table S11**). These results demonstrate that target genes of T2D islet enhancer signals are involved in protein transport and secretion.

Among novel target genes, *IGF2BP2* has a strong islet eQTL with T2D enhancer variants and has no known role in T2D-relevant islet biology. As T2D risk alleles are correlated with reduced *IGF2BP2* expression and reduced insulin secretion phenotypes^22^, we hypothesized that reduced *IGF2BP2* expression in islets would increase T2D risk. We thus determined the effects of reduced *IGF2BP2* on islet function using a mouse model. *IGF2BP2/Imp2* is widely expressed in adult mouse tissues including fat, muscle, liver and pancreas^24^, and in the pancreas *Imp2* expression localized to islets and overlapped insulin (**Figure 5A**). We inactivated *Imp2* in mouse beta cells by recombining the *Imp2^flox(f)^* allele with Cre recombinase driven by the rat insulin 2 promoter (*RIP2*-*Cre*) (**Figure S5A**). Immunoblot analysis of extracts from isolated *Imp2^ff^/RIP2*-*Cre* islets confirmed reduced Imp2 abundance compared to *Imp2^ff^* islets (Figure 5B). *Imp2^ff^/RIP2*-*Cre* mice exhibited no overt phenotype and gained weight similar to *Imp2^ff^* controls on both a normal chow (NCD) and high fat diet (HFD) (**Figure S5B**).

**Figure 5.**
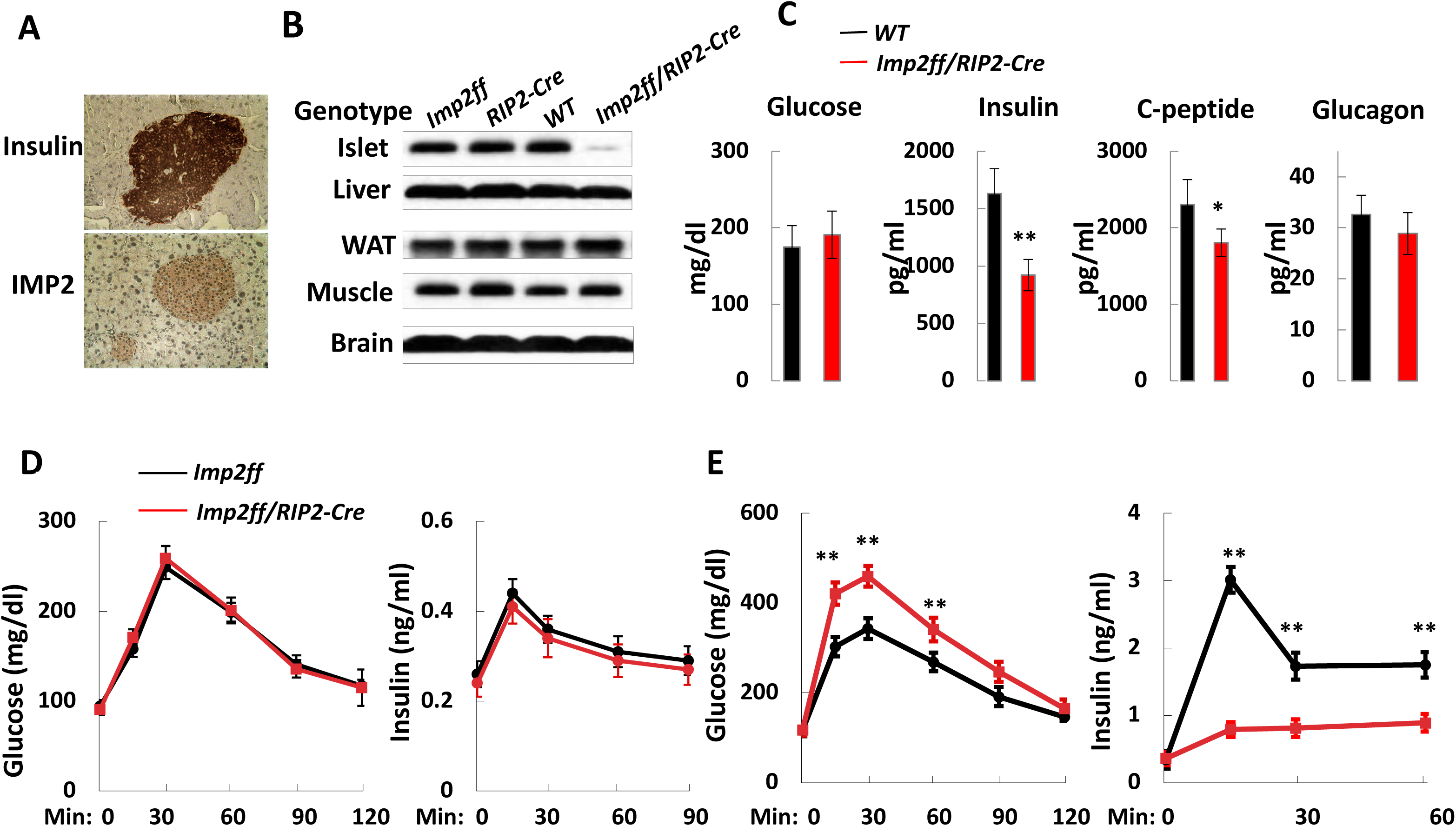
Reduced *IGF2BP2* activity in mouse islets impairs glucose-stimulated insulin secretion during insulin resistance. (A) Immunostaining of insulin and IMP2 in mouse pancreas. (B) Expression of IMP2 in islets and other T2D-relevant tissues liver, adipose, muscle, and brain. (C) Blood glucose, insulin, c-peptide and glucagon level in 10-week-old male mice on high fat diet (HFD) (N=9). Wild-type (black) and *Imp2ff/RIP2*-*Cre* (red). (D) 1 g/kg glucose was administered intraperitoneally after overnight fasting of 12-week-old *Imp2ff* (black; N=10) and *Imp2ff/RIP2*-*Cre* (red; N=10) male mice maintained on normal chow diet (NCD). left=blood glucose; right=serum insulin. (E) 1 g/kg glucose was administered intraperitoneally after overnight fasting to 12-week-old *Imp2ff* (black; N=9) and *Imp2ff/RIP2*-*Cre* (red; N=9) male mice maintained on NCD. left=blood glucose; right=serum insulin. Values represent mean and SD. ^⋆⋆^P<.05, ^⋆⋆^P<.01

We next assessed the effect of *IGF2BP2* deficiency in mouse beta cells on glucose homeostasis. At 10 weeks of age, *Imp2^ff^* and *Imp2^ff^/RIP2*-*Cre* mice on NCD exhibited no difference in blood glucose and insulin levels. By contrast, blood insulin and C-peptide levels were reduced in HFD-fed *Imp2^ff^/RIP2*-*Cre* compared to HFD-fed control mice, whereas blood glucose and glucagon levels were similar (**Figure 5C**). When challenged with an intraperitoneal glucose injection, HFD-fed, but not NCD-fed, *Imp2^ff^/RIP2*-*Cre* mice exhibited significantly higher glucose and lower insulin levels than *Imp2^ff^* mice (**Figure 5D,E**). Importantly, this was not due to a difference in insulin sensitivity, as blood glucose levels after an intraperitoneal insulin injection were similar in *Imp2^ff^* and *Imp2^ff^/RIP2*-*Cre* mice (**Figure S5C**). These results indicate that *IGF2BP2* deficiency limits the capacity of beta cells to augment insulin secretion in response to increased insulin demand.

In summary, we defined the genomic location, function, and spatial orientation of regulatory elements in pancreatic islets. Islet active regulatory elements preferentially interacted with other active elements, in many cases at distances over 1MB, and we identified putative target genes for thousands of islet distal enhancers. Target genes of T2D islet enhancer signals were specifically involved in processes related to protein transport and secretion, and we validated that reduced activity of a previously unknown target gene *IGF2BP2* in mouse islets leads to defects in glucose-stimulated insulin secretion. Together our results define distal regulatory networks in islets and link T2D risk to enhancer regulation of protein transport and secretion. Furthermore, these results highlight the utility of high-resolution chromatin conformation maps in dissecting the gene regulatory networks underlying genetic risk of T2D and other complex disease.

## Methods

### Islet ATAC-seq data generation

Four human islet donors were obtained from the Integrated Islet Distribution Program (IIDP) (**Table S1**). Islet preparations were further enriched and selected using zinc-dithizone staining. We generated ATAC-seq data from the four human islet samples with a protocol as previously described^11^. For each sample, we trimmed adaptor sequences using TrimGalore (https://github.com/FelixKrueger/TrimGalore). The resulting sequences were aligned to sex-specific hg19 reference genomes using bwa mem^25,26^. We filtered reads were to retain those in proper pairs and with mapping quality score greater than 30. We then removed duplicate and non-autosomal reads. We called peaks individually for each sample with MACS2^12^ at a q-value threshold of .05 with the following options “—no-model”, “—shift -100”, “—extsize 200”. We removed peaks that overlapped genomic regions blacklisted by the ENCODE consortium and merged the peaks^26^. In total, we obtained 105,734 merged peaks. To assess concordance between ATAC-seq experiments, we calculated read coverage at 200 bp bins genome-wide, excluding blacklisted genomic regions. We then calculated the Pearson correlation between the read counts for each sample.

### Islet Hi-C data generation

We generated Hi-C data from three pancreatic islet samples, two of which also had ATAC-seq data (**Table S1**). Islet preparations were further enriched and selected using zinc-dithizone staining. *In situ* Hi-C was performed using a previously published protocol with modifications adapted to frozen human tissue^17^. Briefly, the tissue was cut to fine pieces and washed with cold PBS. Cross-linking was carried out with 1% formaldehyde (sigma) in PBS at room temperature (RT) for 10 min and quenched with 125mM Glycine (sigma) at RT for 5 min. Nuclei were isolated using a loose-fitting douncer in hypotonic buffer (20mM Hepes pH7.9, 10mM KCl, 1mM EDTA, 10% Glycerol and 1mM DTT with additional protease inhibitor (Roche) for 30 strokes and centrifuge at 4 °C.

Nuclei were digested using 4 cutter restriction enzyme MboI (NEB) at 37 °C overnight (o/n). Digested ends were filled in blunt with dBTP with biotinylated-14-ATP (Life Technologies) using Klenow DNA polymerase (NEB). Re-ligation was performed *in situ* when nucleus was intact using T4 DNA ligase (NEB) at 16 °C for 4 hrs. The cross-linking was reversed at 68 °C o/n while protein was degraded with proteinase K treatment (NEB). DNA was purified with phenol-chloroform extraction and ethanol precipitation, followed by fragmentation to 300-500 bp with the Covaris S220 ultrasonicator. Ligation products were enriched with Dynabeads My One T1 Streptavidin beads (Life Technologies). PCR was used to amplify the enriched DNA for sequencing. HiSeq 4000 sequencer (Illumina) was used to sequence the library with 2×100 bp paired-end reads.

For each sample, reads from paired end reads were aligned with bwa mem^27^ as single-end reads, and then filtered through following steps. First, only five prime ends were kept for chimeric reads. Second, reads with low mapping quality (<10) were removed. Third, read ends were then manually paired, and PCR duplicates were removed using Picard tools (https://github.com/broadinstitute/picard). Finally, filtered contacts were used to create chromatin contact maps with Juicebox^28^.

Contact maps for each sample were binned to 100kb, and the correlation between samples across all bins for all chromosomes was calculated using scipy.stats.pearsonr in scipy. Chromatin loops were identified by using HICCUPS^8^ at 5kb, 10kb, and 25kb resolutions with default parameters on the Hi-C maps for each individual. The Hi-C data was then pooled across all three samples to create a single contact map, and loops were called with HICCUPs at the same resolutions with the same parameters. A single loop set was then created by identifying loops where both anchors were within 20kb of one another via pgltools^29^ and retaining the loop with the highest resolution. If multiple loops were found at the highest resolution, loops were kept from the contact map with the highest overall sequencing depth. We also called topologically associated domains (TADs) from the pooled Hi-C data using a previously described approach^10^.

### Islet ChIP-seq data processing

We obtained previously published data from ChIP-seq assays of H3K4me1, H3K27ac, H3K4me3, H3K36me3 and CTCF generated in primary islets and for which there was matching input sequence from the same sample^4–6^. For each assay and input, we aligned reads to the human genome hg19 using bwa^30^ with a flag to trim reads at a quality threshold of less than 15. We converted the alignments to bam format and sorted the bam files. We then removed duplicate reads, and further filtered reads that had a mapping quality score below 30. Multiple sequence datasets obtained from the same assay in the same sample were then pooled.

We defined chromatin states from ChIP-seq data using ChromHMM^14^ with a 9-state model, as calling larger state numbers did not empirically appear to identify additional states. We assigned the resulting states names based on patterns previously described in the NIH Roadmap and ENCODE projects – CTCF (CTCF), Transcribed (Txn; H3K36me3), Active promoter (TssA; H3K4me3, H3K4me1), Flanking promoter (TssFlnk; H3K4me3, H3K4me1, H3K27ac), Weak/Poised Enhancer (EnhWk; H3K4me1), Active Enhancer 1 (EnhA-1; H3K27ac), Active Enhancer 2 (EnhA-2; H3K27ac, H3K4me1), and two Quiescent states (Quies; low signal for all assays).

We then annotated accessible chromatin sites based on overlap with the chromatin states. If an accessible chromatin site overlapped multiple chromatin states, we split the site into multiple distinct elements.

### Islet chromatin interaction analyses

To determine the normalized tag counts of ATAC-seq data at loop anchors, loop anchors were converted to a regular BED file with pgltools^29^, and HOMER^31^ was used to find the normalized tag density across all loop anchors for each ATAC-seq sample. Output from HOMER was normalized to a maximum height of 1 for each sample to determine the distribution of ATAC-seq signal within each sample, rather than the relative magnitude coverage difference between ATAC-seq samples.

To determine the enrichment of each class of islet regulatory elements near loops, and the types of elements colocalized by loops, we utilized pgltools and HOMER to integrate the ATAC-seq and Hi-C data. We first created a size matched null distribution comprised of 7,000 permuted regions. Next, for each islet accessible chromatin state, we identified the proportion of states within 25kb of a loop. We determined the fold enrichment of each class over the average calculated from the null distribution, and determined significance as the number of permuted counts greater than the observed.

To determine which pairs of islet regulatory elements were in chromatin loops at a statistically significant level, we compared the distribution of islet regulatory elements around loop anchors using HOMER. We utilized the “annotateInteractions” function to obtain logistic regression p-values and odds ratio enrichment estimates for all pairs of islet regulatory elements.

We defined candidate target genes of islet enhancer elements using Hi-C loops in the following way. First, we identified all islet active enhancer elements mapping within 25kb of a Hi-C loop anchor. We then filtered these loops based on whether the other anchor mapped within 25kb of a promoter region (-5kb/+2kb of transcription start site) for protein-coding or long non-coding genes in GENCODEv27^32^. For each active enhancer, we then calculated the number of gene promoter regions interacting with that enhancer. For each gene promoter region, we calculated the number of independent interactions containing at least one active enhancer element.

We identified genes with multiple (>2) active enhancer interactions and tested these genes for gene set enrichment using GSEA^33^, considering only gene sets with more than 25 genes at an FDR>.2.

### Genomic enrichment analyses

We tested for enrichment of variants in each accessible chromatin class using T2D association data of 1000 Genomes project variants from the DIAGRAM consortium^21^. From this meta-analysis, we identified common variants (with minor allele frequency (MAF)>.05). In total, we retained 6.1M common variants for testing. For each variant, we then calculated a Bayes Factor from effect size estimates and standard errors using the approach of Wakefield^34^.

We then modelled the effect of variants in each class of islet regulatory elements on T2D risk using fgwas^20^. For these analyses, we used a window size (-k) that resulted in a 1Mb window on average. We first tested for enrichment of variants in each state individually. We then built a joint model iteratively in the following way. We first identified the annotation with the highest likelihood. We then added annotations to the model until the likelihood did not increase further. Using this model, we introduced a series of penalties from 0 to .5 in increments of .01 and fit the model using each penalty, and identified the penalty that gave the highest cross-validation likelihood. We then finally removed annotations from the model that further increased the cross-validation likelihood. We considered the resulting set of annotations and effects to be the optimal joint model.

We also modelled the effect of variants in islet regulatory elements using LD-score regression. For these analyses, we extracted variants in HapMap3 from T2D association data. We then calculated LD scores for variants in each regulatory element class. Finally, we obtained enrichment estimates using these LD scores with T2D association data of HapMap3 variants.

### Fine-mapping of T2D risk variants

We used the effects from the joint enrichment model as priors on the evidence for variants at 107 known T2D signals using fine-mapping data from the Metabochip^2^, GoT2D^1^ and DIAGRAM 1000 Genomes^21^ studies. We used data of 49 T2D signals at 39 T2D loci on the Metabochip, 41 additional T2D signals from GoT2D data for T2D loci not on the Metabochip, and 17 additional T2D signals in DIAGRAM 1000G not in Metabochip or GoT2D.

For each signal, we obtained the enrichment effect of the islet regulatory or coding annotation overlapping that variant. We calculated a prior probability for the variant by dividing the effect by the sum of effects across all variants at a signal. We then multiplied this prior probability by the Bayes Factor for each variant. From the resulting odds, we calculated a posterior probability that the variant is causal for T2D risk (PPA) by dividing the odds by the sum of odds across all variants at the locus.

For each signal, we calculated a cumulative PPA (cPPA) value for islet enhancer (EnhA1, EnhA2, EnhWk), promoter (TssA, TssFlnk), CTCF binding site, UTR, and coding exon (CDS) annotations by summing the PPAs of all variants overlapping each annotation. We then clustered T2D signals into groups based on cPPA values using k-means clustering.

We determined the effects of T2D signals in each cluster on glycemic association data^22^. We identified 73 T2D signals represented in these data and cataloged 23 associated at P<.05 with first-phase insulin response, peak insulin response, AIR, or insulin secretion rate. We calculated the percentage of signals in each cluster associated with these measures and tested for differences between clusters using a chi-square test.

For the 30 T2D islet enhancer signals, we calculated “99% credible sets” as the set of candidate variants that explain 99% of the total PPA using genetic fine-mapping data alone (genetic), and fine-mapping including priors from the joint genome-wide enrichment model (+priors).

We then fine-mapped casual variants in putative T2D loci genome-wide. For variants in each 1MB window across the genome, after excluding any windows overlapping a known T2D signal, we obtained the effect of the islet annotation overlapping that variant. We calculated a prior probability for each variant as described above also including an additional prior on the evidence that the 1MB window is a T2D locus. We multiplied both prior probabilities by the Bayes Factor for each variant. From the resulting odds, we calculated the PPA that each variant is causal for T2D risk. We then considered the 131 windows with at least one islet enhancer variant with PPA>.01 in downstream analyses.

### Genomic features analyses

For each class of islet open chromatin, we determined the overlap with other genomic features.

We identified motifs enriched in sequence underneath each islet accessible chromatin class. We first extracted genomic sequence for each site using bedtools^35^, and masked repetitive sequences. We then identified *de novo* motifs enriched in this sequence using DREME^36^. For each *de novo* motif, we determined whether this motif matched a known sequence motif in a custom database of >2,500 motifs from ENCODE, JASPAR and SELEX with tomtom^26,37–39^.

We determined the overlap of islet accessible chromatin classes with transcription factor (TF) ChIP-seq data in islets for 5 proteins^4,26^. For each islet chromatin class, we calculated the Jaccard index of overlap with sites for each TF^35^. We then determined the overlap of islet accessible chromatin classes with DHS sites from 384 cell types in the ENCODE project^26^. We first filtered out DHS sites from islets, and then for each accessible chromatin site, we calculated the percentage of ENCODE cell-types the site was active in. We then determined the median percent overlap across all sites within each accessible chromatin class.

### Gene expression analysis

We obtained transcript-per-million (TPM) counts from RNA-seq data in 53 tissues from the GTEx project release v7^19^. We further obtained RPKM read counts from RNA-seq data of 118 pancreatic islet samples^18^, and calculated TPM values as previously described^40^. We then retained only protein-coding and long non-coding genes annotated in GENCODEv27^32^. We first calculated the percentage of genes expressed in islets (defined as ln(TPM)>1) and not expressed in islets with at least one enhancer chromatin loop to the promoter region and tested for a significant difference using a Chi-square test.

Across all 54 tissues, we filtered out genes not expressed (ln(TPM)>1) in at least one tissue. We determined correlation between gene expression level in islets and enhancer loop number using Spearman’s rho. We further grouped genes by the number of chromatin loops to enhancer elements and calculated the median islet TPM value for each group. For genes with at least one enhancer loop, we created a linear model of log-transformed gene TPM with chromatin loop number as the predictor using the glm package in R. Values are reported as the p-value, effect size (beta) and standard error from the resulting model.

We then determined the relative expression level for each gene in 54 tissues. We log-transformed expression values and calculated a z-score for each gene using the mean and standard deviation across tissues. We then repeated the above analyses using tissue z-scores instead of tissue TPM values.

### Islet expression QTL analysis

We obtained summary statistic eQTL data from two published studies of 118 and 112 primary pancreatic islet samples^18,41^. We then performed inverse sample-size weighted meta-analysis to combine the summary results for each variant and gene pair using METAL^42^. We retained only protein-coding and long non-coding RNA genes as defined by GENCODEv27^32^, only variant and gene pairs tested in both studies, and only variants with minor allele frequency (MAF) > .01.

We extracted eQTL associations for variants in classes of islet accessible chromatin. To remove potential biases due to linkage disequilibrium, we sorted variant associations based on p-value and iteratively pruned out variants in LD (r^2^>.5) with a more significant variant using LD information in European samples from 1000 Genomes project data. We then extracted pruned eQTL associations for variants in active promoter elements for genes within 20 kb (TssA), variants in active enhancer elements for genes within 20 kb (EnhA proximal), variants in active enhancer elements for genes in chromatin loops (EnhA distal target), and variants in active enhancer elements for genes without a loop (EnhA distal no-target). For each set of eQTL associations, we compared p-value distributions using a two-sided Wilcox rank-sum test. To remove potential biases in variant distances to loop and no-loop genes, we randomly selected variant-gene pairs matched on distance to the distal target set to re-performed analyses.

### Target genes of T2D islet enhancer signals

We defined candidate target genes of 30 known T2D enhancer signals and 131 putative T2D enhancer windows in the following way. We identified candidate causal variants at each signal overlapping islet enhancer elements and considered target genes as those where a candidate variant either (a) mapped in a chromatin loop to the promoter region (-5kb/+2kb of transcription start site) or (b) was within 25kb proximal to the promoter region.

We next defined alternate sets of target genes of the 30 T2D enhancer signals based on 1MB windows or TAD boundaries. For 1MB window definitions, we identified the highest probability variant for each signal and extracted a +/-1MB window around the variant position. We then considered gene promoter regions (-5kb/+2kb of transcription start site) for protein-coding or long non-coding genes in GENCODEv27 that overlapped this +/-1MB window the set of target genes. For TAD boundary definitions, we intersected the merged set of TADs with gene promoter regions to obtain a set of genes within each TAD. We then intersected the highest probability variant at each T2D signal with TADs to obtain gene sets in the TAD.

For each enhancer signal with a candidate target gene, we extracted eQTL p-values for each target gene using the islet enhancer variant with the highest PPA at the signal. Where the highest probability variant was not present in the eQTL dataset, we used the next most probable islet enhancer variant. We then Bonferroni-corrected eQTL p-values for the total number of candidate target genes at the signal and considered eQTLs significant with a corrected p-value<.05.

For genes with significant eQTL evidence we further tested whether T2D and eQTL signals were co-localized. We obtained the T2D Bayes Factor for each variant at the signal from fine-mapping data. For significant gene eQTLs at the signal, we then calculated the Bayes Factor that each variant is an islet eQTL for that gene^34^. We compared Bayes Factors for T2D signals and eQTLs for each gene using Bayesian co-localization^43^. We considered the prior probability that a variant was causal for T2D risk or an islet eQTL as 1×10^−4^, and the prior probability that a variant was causal for both T2D risk and an islet eQTL as 1×10^−5^. We considered signals as having evidence for co-localization where the probability of a shared causal variant was higher than the probability of two distinct causal variants.

We tested target genes for gene set enrichment using GSEA^33^, considering only gene sets with more than 25 total genes and at an FDR threshold of .2.

### Luciferase reporter assays

To test for allelic differences in enhancer activity at rs7732130, we cloned sequences containing alt or ref alleles in forward and reverse orientation upstream of the minimal promoter of firefly luciferase vector pGL4.23 (Promega) using KpnI and SacI restriction sites.

The primer sequences were:

forward/left: GATAACGGTACCGCGAAGTGGTCATGGGTAAA
forward/right: AAGTAGGAGCTCACCATCCCAGCATTTAGTGG
reverse/left: GATAACGAGCTCGCGAAGTGGTCATGGGTAAA
reverse/right: AAGTAGGGT ACCACCATCCCAGCATTTAGTGG

MIN6 beta cells were seeded into 6 (or 12)-well trays at 1 million cells per well. At 80% confluency, cells were co-transfected with 400ng of the experimental firefly luciferase vector pGL4.23 containing the alt or ref allele in either orientation or an empty vector and 50ng of the vector pRL-SV40 (Promega) using the Lipofectamine 3000 reagent. All transfections were done in triplicate. Cells were lysed 48 hours after transfection and assayed for Firefly and Renilla luciferase activities using the Dual-Luciferase Reporter system (Promega). Firefly activity was normalized to Renilla activity and compared to the empty vector and normalized results were expressed as fold change compared to empty vector control per allele. Error bars are reported as standard deviation. A two-sided t-test was used to compare luciferase activity between the two alleles in each orientation.

### Mouse Imp2 targeting construct and physiological studies

We generated the Imp2 construct by using a genomic fragment of 12 kb containing Imp2 exons 1 and 2 as well as flanking intron sequences of the murine gene extracted from the RP23-163F16 BAC clone. The replacement-type targeting construct consisted of 9.4 kb of Imp2 genomic sequences (4.4 kb in the left homology arm and 5.4 kb in the right homology arm) (**Figure S5**).

We bred mice for experiments by crossing IMP2-loxp mice (*Imp2^ff^*) with RIP2-Cre mice on a C57Bl/6J background. We maintained colonies in a specific pathogen-free facility with a 12:12 light - dark cycle and fed irradiated chow (Prolab 5P75 Isopro 3000; 5% crude fat; PMI nutrition international) or a HFD (D12492i; 60% kcal fat; Research Diets Inc.). Blood glucose, insulin, C-peptide and glucagon levels were measured by the Vanderbilt metabolic core. Measurements for *Imp2^ff^* and *Imp2^ff^/RIP2*-*Cre* mice were performed using male mice under basal conditions (N=10), upon intraperitoneal glucose injection (N=9), and upon intraperitoneal insulin injection (N=9). A two-sided t-test was used to compare differences in measurements across genotypes.

## Acknowledgements

Support for this work was provided by U01DK105541 to M.S., K.F. and B.R., the Ludwig Institute for Cancer Research to B.R., R37DK017776 and P30DK057521 to J.A., postdoc fellowship 537-2014-6796 from the Swedish Vetenskapsrådet to J.Y., and JDRF-3-2012-177 postdoc fellowship to A.W.

## Author Contributions

K.J.G., B.R., M.S., K.F. conceived of and supervised the research in the study; K.J.G. wrote the manuscript and performed analyses; W.W.G, J.C., Y.Q. performed analyses and contributed to writing; J.Y. performed Hi-C assays and contributed to writing; N.D., J.A. performed mouse experiments and contributed to writing; A.W., A.A. contributed analyses and data interpretation; J.Y.H., N.V., F.D., D.G. performed ATAC-seq assays and contributed to data interpretation; N.K. and M.O. performed variant reporter experiments; L.B. and L.M. contributed to mouse experiments.

## Data availability

Data files for this study are available at http://gaultonlab.org/pages/Greenwald_islet_HiC and will also be deposited in https://www.t2depigenome.org

## Supplemental Figures

**Figure S1.**
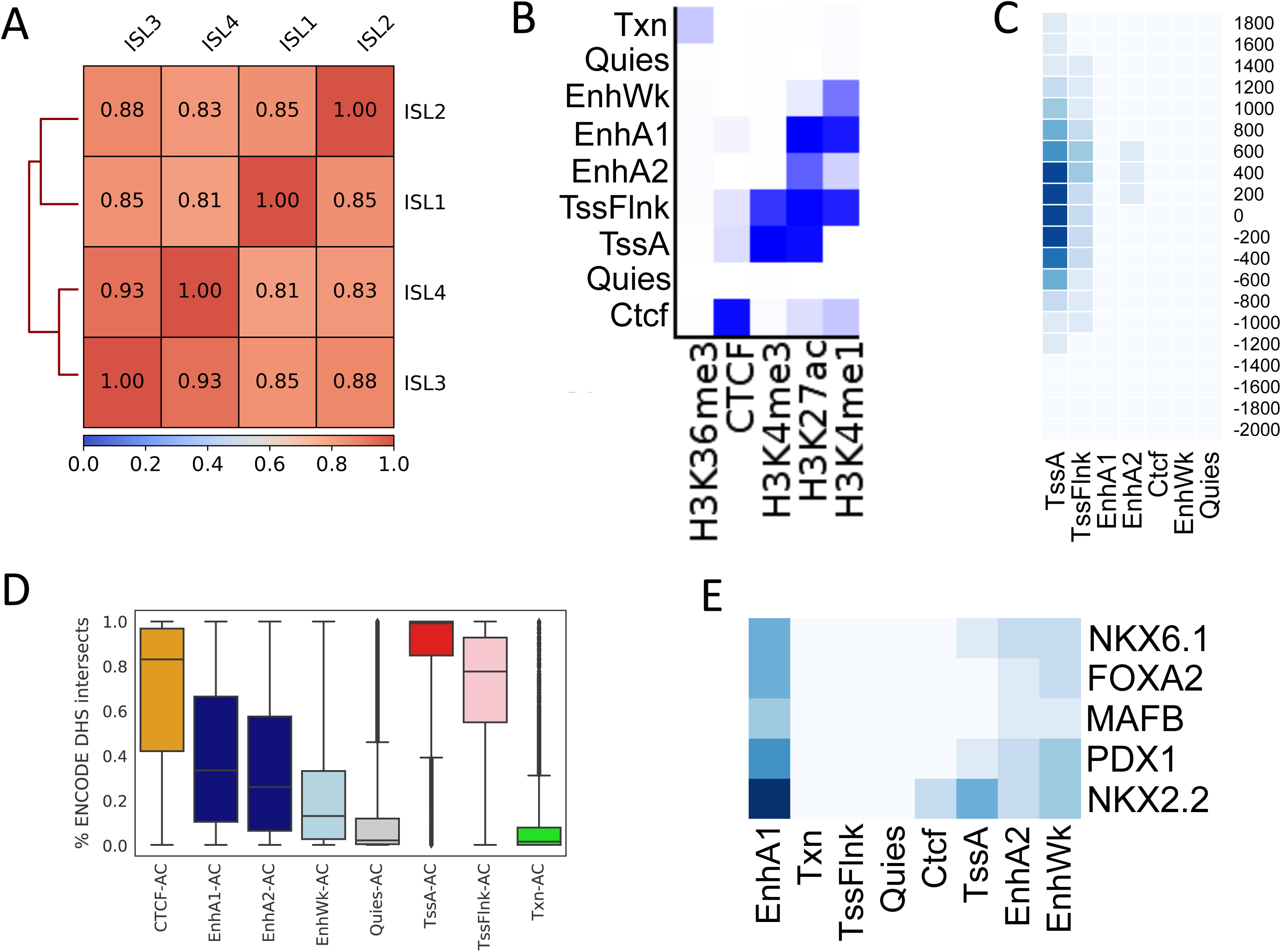
Characteristics of pancreatic islet regulatory elements. (A) Heatmap of the Pearson correlation of ATAC seq signal across four islet samples, calculated as the raw tag count in 1kb bins across the genome. (B) Heatmap of emission matrix probabilities for the 9-state islet model from chromHMM, with individual ChIP-seq assays shown on the x-axis and labelled chromatin states on the y-axis. (C) Heatmap showing percentage of each class of islet regulatory elements mapping in 200bp bins around GENCODE transcriptional start sites. (D) Percentage of ENCODE cell-types in DHS sites overlapping each class of islet regulatory elements. (E) Jaccard overlap of each class of islet regulatory elements with islet ChIP-seq sites for five transcription factors.

**Figure S2.**
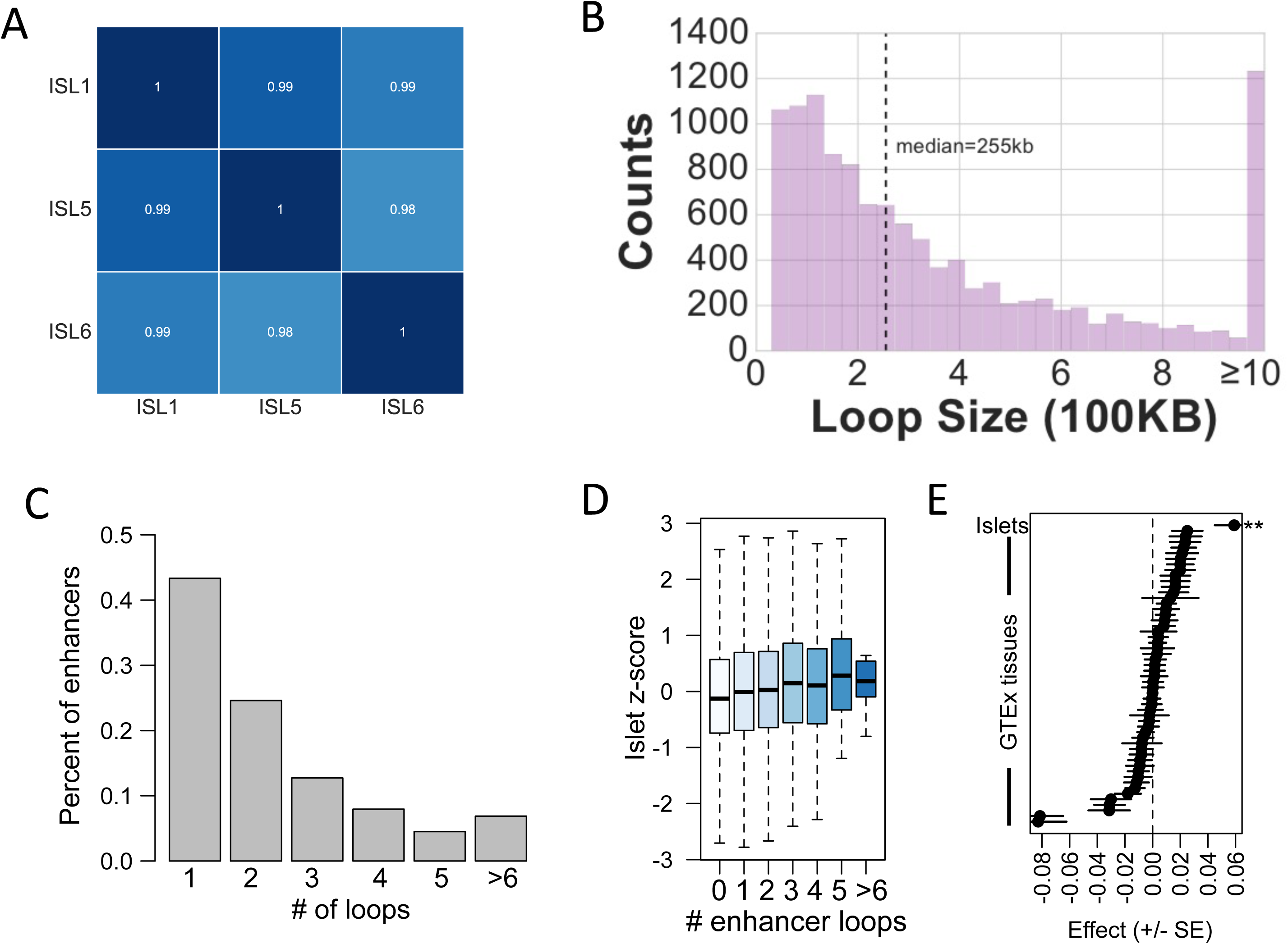
Characteristics of pancreatic islet chromatin loops. (A) Heatmap showing the Pearson correlation of Hi-C contacts across islet samples in 100kb bins across the genome. (B) Histogram of the distance between loop anchors in islets. (C) Histogram of number of loops within 25kb of each islet active enhancer to gene promoter regions. (D) Boxplot showing genes with increasing numbers of chromatin loops to islet enhancers (x-axis) had on average higher relative expression level in islets (y-axis). (E) The number of chromatin loops to islet enhancers was a significant predictor of relative gene expression level in islets but not 53 other tissues in GTEx. ^⋆⋆^P<.001. Values represent effect size and SE.

**Figure S3.**
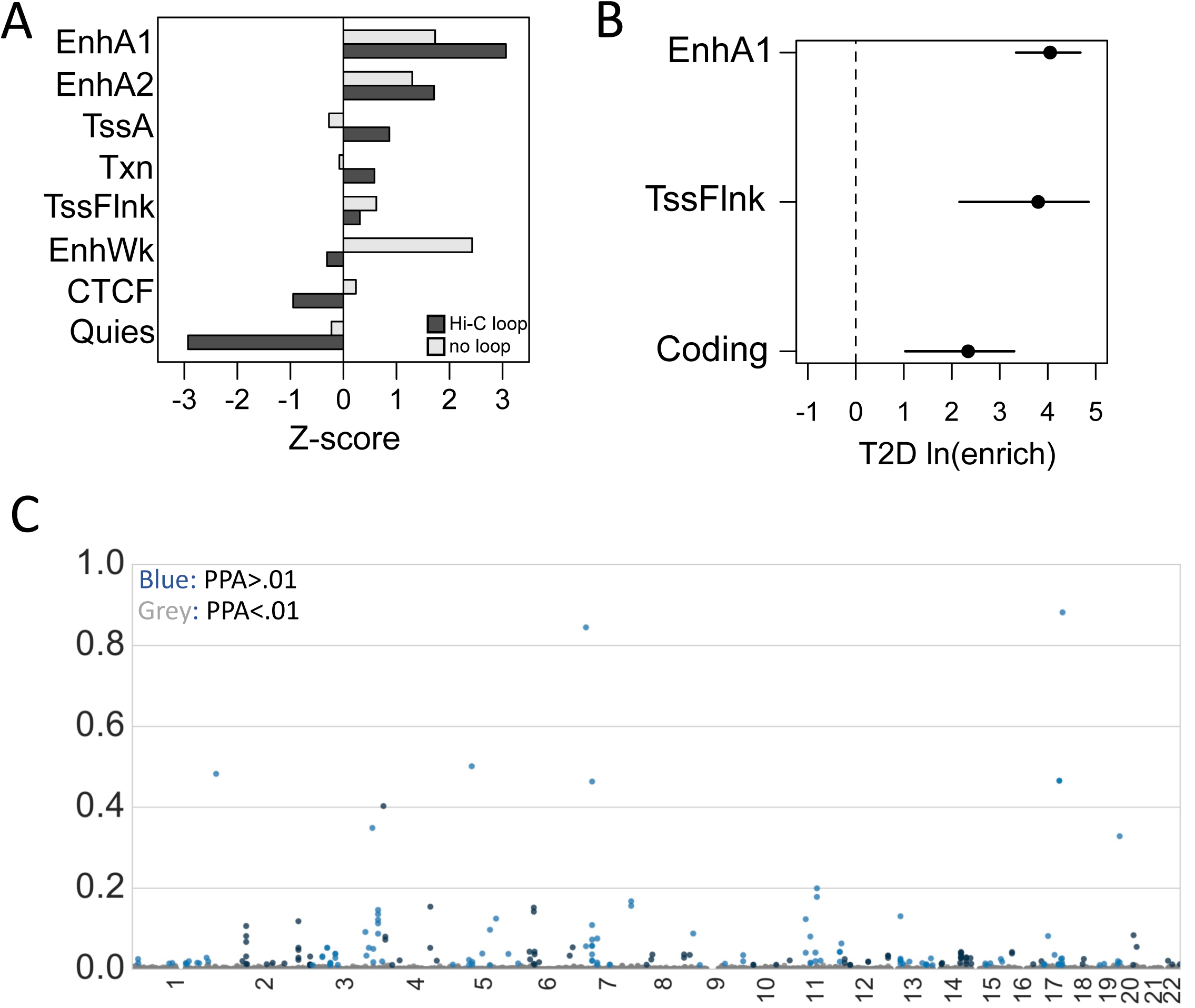
Effects of variants in pancreatic islet regulatory elements on T2D risk. (A) Enrichment Z-score measured using LD-score regression for each class of islet regulatory elements (y-axis), subset by states that were (dark) or were not (light) within 25kb of a Hi-C loop anchor. (B) Enrichments from the fgwas genome-wide joint model including islet active enhancers (EnhA1), flanking promoters (TssFlnk), and coding exons (CDS). Values represent log enrichment and 95% CI. (C) Posterior causal probabilities (PPA) of variants within islet active enhancers in 1MB windows genome-wide excluding known T2D risk loci. Blue = variants with PPA>0.01, grey = variants with PPA<0.01.

**Figure S4.**
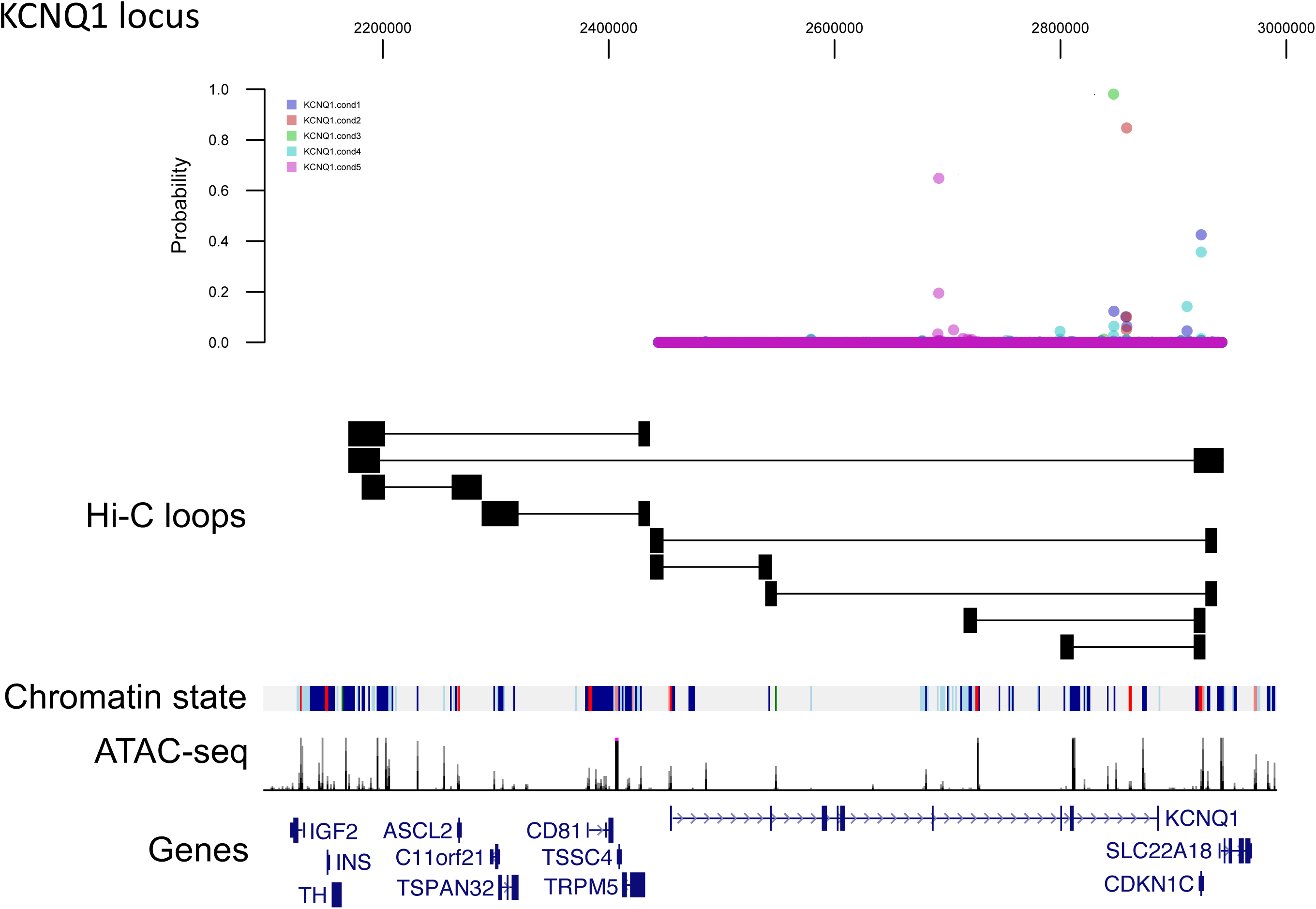
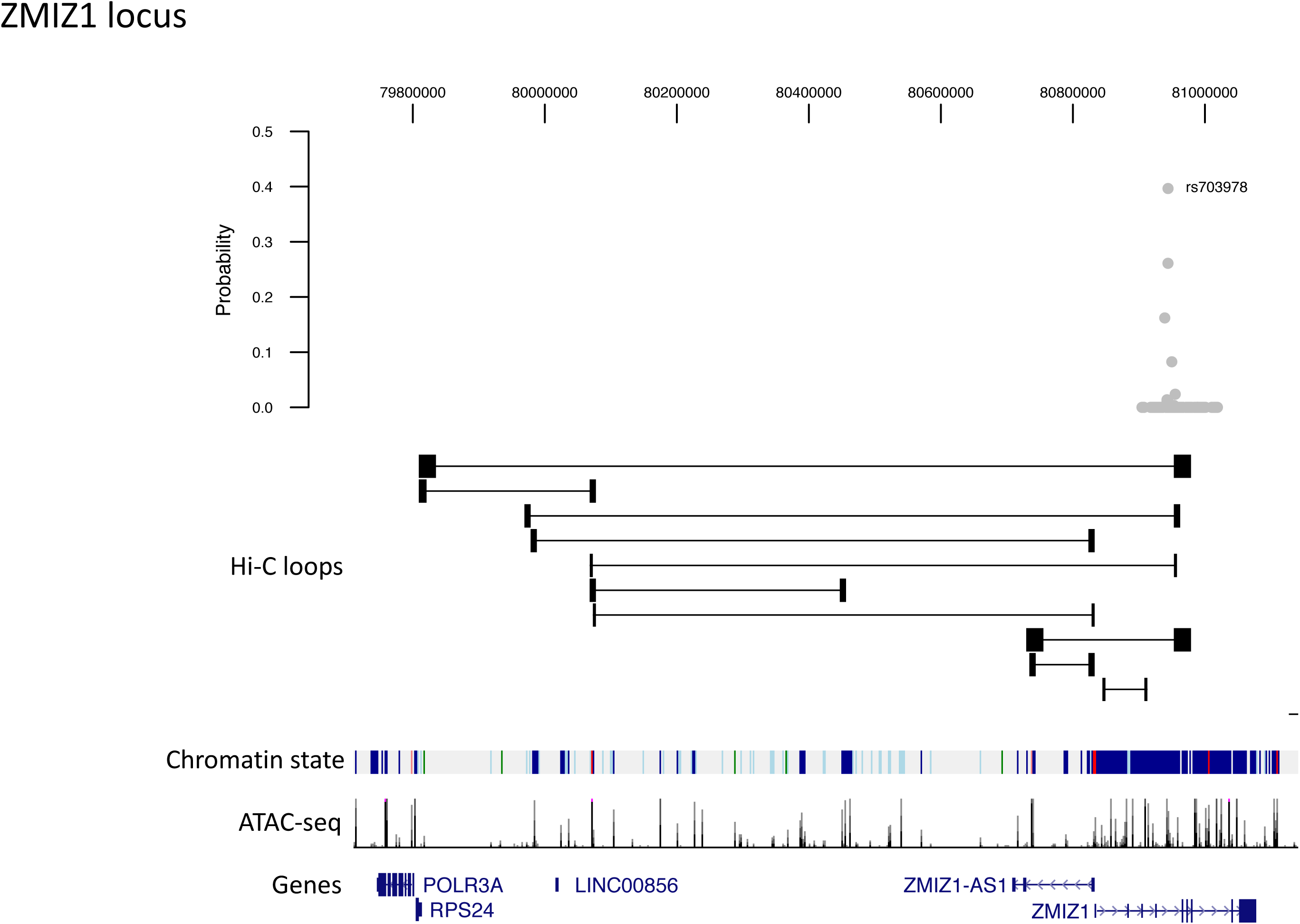
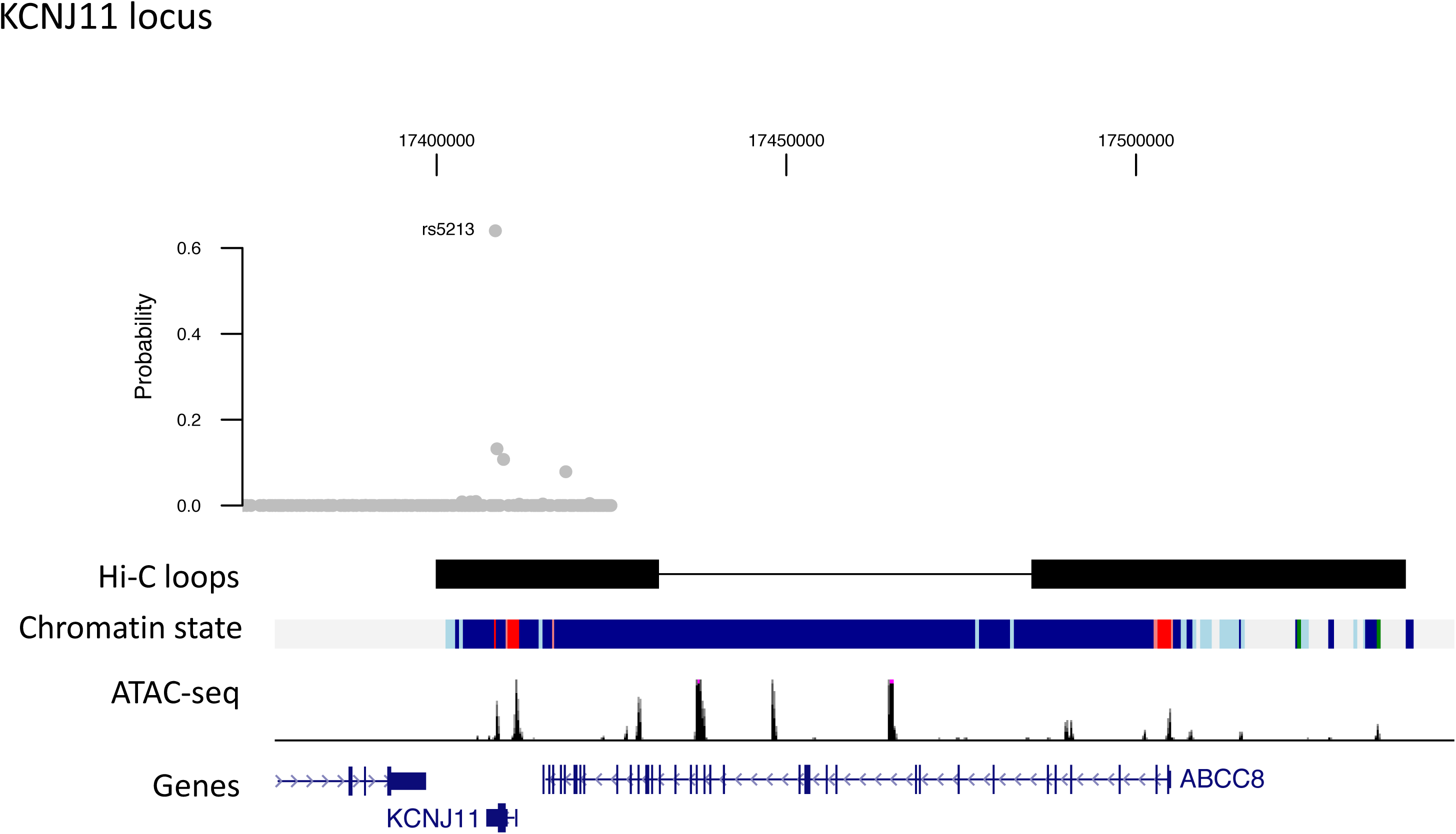
T2D enhancer signal chromatin loops to candidate target genes. Re-weighted posterior causal probabilities of variants (top), islet Hi-C loops, chromatin states and ATAC-seq signal (middle), and known genes (bottom), for (A) five independent T2D risk signals at the *KCNQ1* locus, (B) T2D signal at the *ZMIZ1* locus, and (C) T2D signal at the *KCNJ11/ABCC8* locus. For (A), posterior probabilities are shown in different colors for each of the five independent signals.

**Figure S5.**
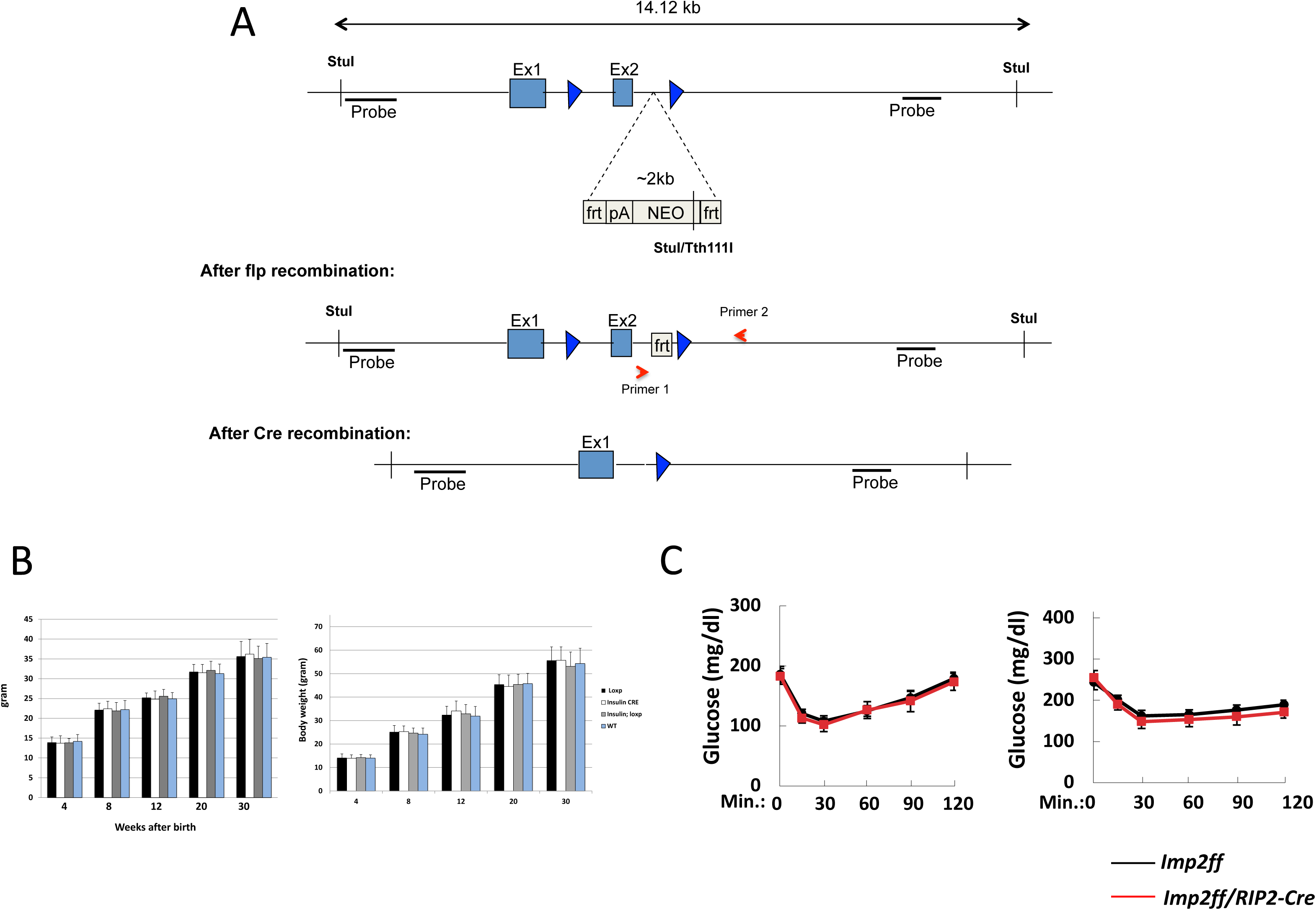
Characterization of mice after conditional *IGF2BP2* ablation in beta cells. (A) Schematic representation of the wild type *Imp2* allele showing exon 1-2 and flanking intron sequences, and the *Imp2*^flox^ targeted allele. (B) Body weight for wild-type, *RIP2*-*Cre*, *Imp2^ff^* and *Imp2^f^/RIP2*-*Cre* mice on normal chow diet (top) and high fat diet (bottom). (C) Insulin tolerance tests in 14-week-old *Imp2^ff^/RIP2*-*Cre* (red) and *Imp2^ff^* (black) mice on a normal chow diet (left) and high fat diet (right). Values represent mean and SD.

## Supplemental Tables

**STable 1:** Donor and sequencing characteristics of pancreatic islet samples

**STable 2:** Regulatory elements in pancreatic islets

**STable 3:** Sequence motifs enriched in islet regulatory elements

**STable 4:** Hi-C loops identified in pancreatic islet samples

**STable 5:** Target gene chromatin loops of islet enhancer elements

**STable 6:** Islet enhancer chromatin loops of gene promoter regions

**STable 7:** Functional annotations enriched in genes with multiple enhancer interactions

**STable 8:** T2D candidate causal variants in islet active enhancers

**STable 9:** Target genes of T2D islet enhancer signals

**STable 10:** Gene set annotations enriched in target genes of T2D islet enhancer signals

**STable 11:** Target genes with islet eQTLs for T2D islet enhancer signals

